# Additive framework of hormonal waves explains species and age differences in circadian intraocular pressure rhythm

**DOI:** 10.1101/2025.10.21.683671

**Authors:** Keisuke Ikegami, Ryo Fujie, Fumito Mori, Shinobu Yasuo

## Abstract

Elevated intraocular pressure (IOP) is the primary risk factor for glaucoma, yet IOP demonstrates significant circadian rhythms, and their disruption heightens disease susceptibility. A paradox exists in that both diurnal and nocturnal animals experience nocturnal IOP elevation despite their contrasting behavioral chronotypes. Here, we developed a minimal mathematical framework where IOP rhythms arise from the linear superposition of two sinusoidal signals: adrenal glucocorticoids (GC) and norepinephrine (NE) from the superior cervical ganglion. In both diurnal and nocturnal species, NE levels increase at night, while GC levels peak oppositely in the morning and evening. A meta-analysis of published datasets showed that IOP peaks in the early night for nocturnal animals and in the late night for diurnal animals, aligning with the predicted maxima of the combined GC and NE sine waves. In aged mice and following superior cervical ganglionectomy, IOP rhythms shifted in phase and decreased in amplitude and mean level; these changes were accounted for by selectively reducing the NE component’s amplitude in the model. Conversely, in diurnal humans, aging results in a delayed IOP phase, which is replicated by diminishing NE amplitude. Thus, species differences, age-related changes, and the effects of sympathetic ablation on IOP can be coherently explained by the combination of the two zeitgeber signals with distinct phases. This straightforward yet robust framework offers a unifying concept for the circadian regulation of IOP across species and may inform the development of novel diagnostic algorithms and chronotherapeutic strategies for glaucoma.

**Significance:** Glaucoma is a leading cause of irreversible blindness, and its major risk factor, intraocular pressure (IOP), exhibits a circadian rhythm. A long-standing paradox is that IOP rises at night in both diurnal humans and nocturnal rodents, despite their opposite activity patterns. We showed that IOP rhythms can be explained by the superposition of two sine waves representing adrenal glucocorticoid and sympathetic norepinephrine rhythms. This framework parsimoniously accounts for species differences, aging effects, and the impact of sympathetic ganglionectomy on the IOP. By reducing a complex physiological process to the interaction of two entrainment signals with distinct phases, our model provides new mechanistic insights into circadian ocular physiology and highlights potential strategies for age-specific monitoring and therapeutic timing in glaucoma.

## Introduction

Glaucoma is the leading cause of irreversible blindness worldwide, and there is no effective cure. Elevated intraocular pressure (IOP) is the most important risk factor; however, IOP fluctuates with a circadian rhythm and a nocturnal peak (1, 2). Patients with glaucoma also exhibit nocturnal IOP elevation, with phase shifts observed in primary open-angle glaucoma and normal-tension glaucoma (NTG)(3). Abnormal IOP rhythms are linked to NTG(4), and aging desynchronizes IOP rhythms in older individuals(5). Disruption of this rhythm in night-shift workers is associated with a higher risk of optic nerve damage and glaucoma(5, 6). These findings highlight the need to regulate nocturnal IOP for effective glaucoma management and underscore the importance of circadian mechanisms. A puzzling feature of IOP regulation is that both diurnal humans (2) and nocturnal rodents (7) exhibit nocturnal IOP elevation despite their opposite activity patterns. The underlying mechanism that generates these convergent rhythms remains unresolved.

Circadian rhythms govern the physiological processes of most organisms. The suprachiasmatic nucleus (SCN) acts as the primary circadian pacemaker, receiving retinal input to regulate biological rhythms(8)including the IOP rhythm(7). The SCN coordinates peripheral tissues via the autonomic nervous system(9). Norepinephrine (NE) from the superior cervical ganglion (SCG) conveys circadian signals to the ciliary body of the eye, affecting pupil size(10). Glucocorticoids (GCs), secreted by the adrenal glands under hypothalamic–pituitary–adrenal axis control, synchronize circadian activity through receptors in peripheral tissues(11). Our studies identified systemic signals influencing IOP rhythm by GCs and NE(12–14). Both exhibit circadian rhythmicity with species-specific phases; however, their combined effects on species differences and aging remain untested.

Here, we propose and validate a minimal mathematical framework in which IOP rhythms emerge from the superposition of two sinusoidal time rhythms: GC and NE. Using published datasets and newly generated experimental data in young and aged mice, as well as after SCG ablation (SCGX), we showed that this model parsimoniously explains species-specific and age-dependent changes of IOP rhythms. We further extended the model to human data, demonstrating that age-related changes in the IOP phase can also be reproduced by attenuating NE amplitude.

## Results

### An additive framework of hormonal waves accounts for the phase differences in IOP rhythm across species

To investigate sine wave integration for constructing the IOP rhythm model, we first reanalyzed our published dataset(31)using a two-way ANOVA with the interaction between ADX and SCGX as factors (Fig. 1A). The analysis revealed significant main effects of both manipulations on the IOP rhythm amplitude. ADX significantly decreased the amplitude (F(1,34) = 22.1, *p* = 4.1 × 10^−5^), and SCGX also significantly decreased amplitude (F(1,34) = 21.7, *p* = 4.8 × 10^−5^). The interaction term (ADX × SCGX) was not significant (F(1,34) = 0.016, *p* = 0.90), suggesting additive rather than synergistic or antagonistic effects (Fig. 1A). To illustrate the factorial design, we plotted the mean IOP rhythm amplitude against the SCGX status, with lines indicating ADX presence or absence. The parallel lines were aligned with statistical insignificance (Fig. 1A), indicating these signals collectively influence circadian IOP rhythm in mice.

**Fig. 1.**
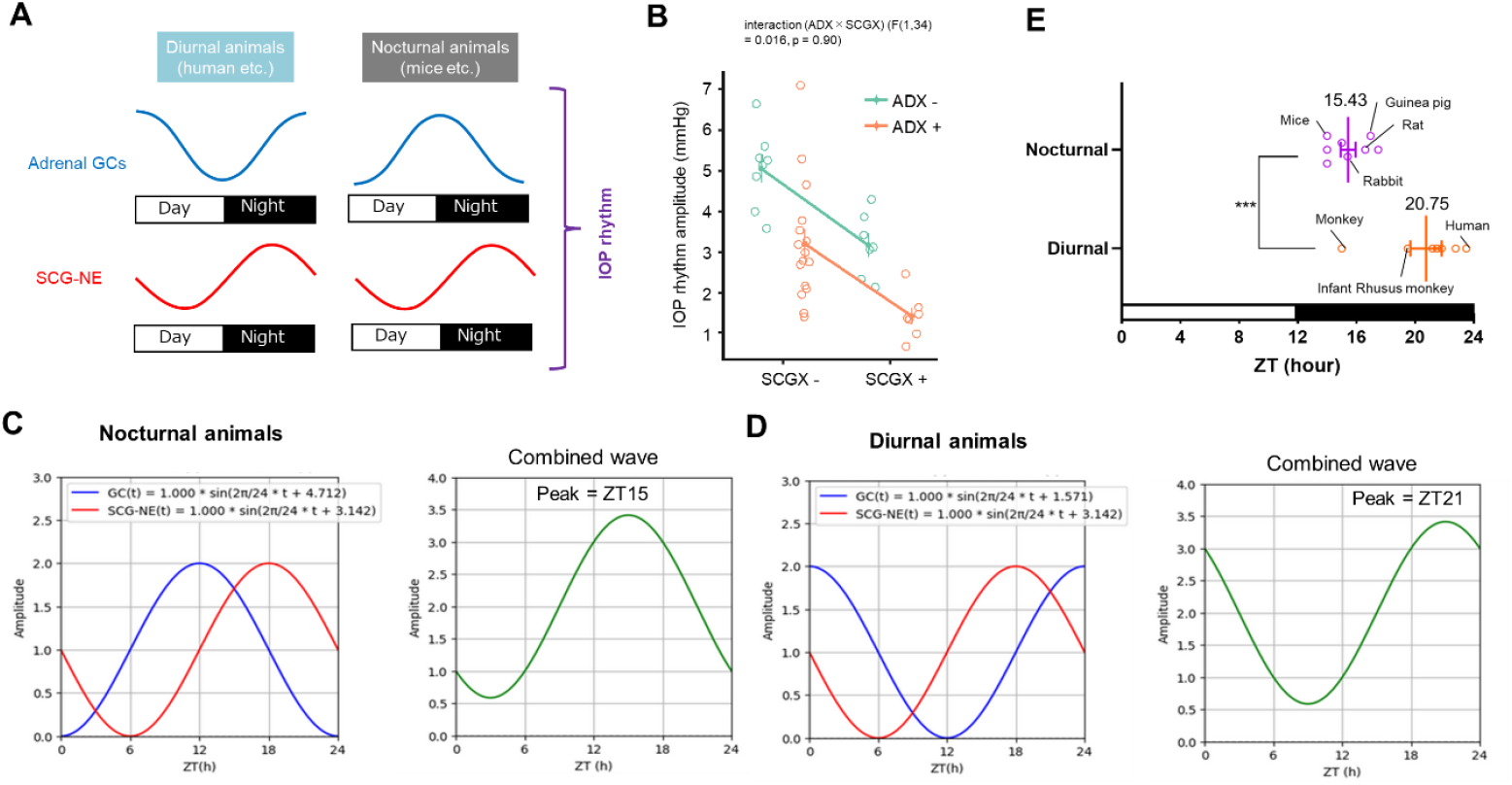
The additive framework of hormonal waves was demonstrated by the phase differences in IOP rhythm observed across species. (A) Conceptual scheme of adrenal glucocorticoid (GC) and superior cervical ganglion norepinephrine (NE) rhythms in diurnal and nocturnal species. GC rhythms are in antiphase, whereas NE rhythms rise during the night in both, suggesting that these two temporal signals jointly shape the IOP rhythms. (B) Scatter plot of IOP rhythm amplitude (reanalysis of our previous IOP rhythm data (31)). Interaction analysis between ADX and SCGX indicated no effect on IOP rhythm amplitude. (C,D) Representative sine waves of GC (blue) and NE (red) in diurnal and nocturnal animals. GC peaks at ZT12 (*a* = −π/2) and NE peaks at ZT18 (*b* = −π), and the peak occurs at *𝜙* =−3π/4, which corresponds to ZT15 in nocturnal animals. In contrast, in diurnal animals, assuming that GC peaks at ZT0 (*a* = π/2) and NE peaks at ZT18 (*b* = −π), the resulting combined wave peaks *𝜙* =−π/4, which corresponds to ZT21. (E) Unweighted meta-analysis of reported IOP rhythm peaks in diurnal (humans and monkeys) and nocturnal (mice, rats, rabbits, and guinea pigs) species. Data are shown as mean ± SEM; scatter points (^***^*p* < 0.001, unpaired t-test).

As we have demonstrated that GC and NE time signals generate IOP rhythm, to explain this conserved nocturnal rise, GC rhythms are in antiphase between diurnal and nocturnal species, whereas SCG-NE consistently rises at night in both species (Fig. 1B). Representative sine waves show this arrangement: NE has a similar nocturnal phase in both chronotypes, whereas GC peaks in the morning for diurnal animals (Fig. 1C), and evening for nocturnal animals (Fig. 1D). Because a normal sine wave oscillates within the ± range, “+*A*” or “+*B*” is added to each sine wave to set the “trough to 0”:

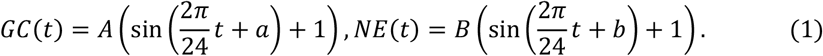

where *t* represents the time (in hours) ranging from 0 to 24, *A* and *B* denote the relative amplitudes (0 ≤ *A, B* ≤ 1), and *a* and *b* represent the phase angles (−π ≤ *a, b* ≤ π). Therefore, to simplify, the combined wave with the same contribution rate is expressed as follows:

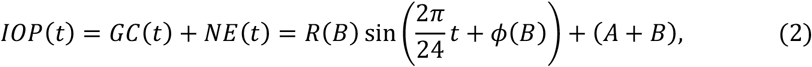

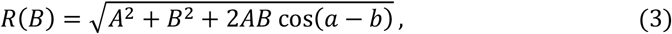

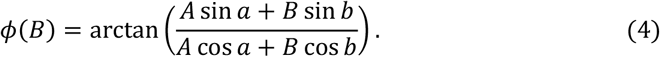

Based on these waveforms, we constructed additive combined sine-wave models of circadian IOP variation (Fig. 1C,D). If we assume that in nocturnal animals, GC peaks at ZT12 (*a* = −π/2) and NE peaks at ZT18 (*b* = −π), with both having an amplitude of 1, and the peak occurs at *𝜙* =−3π/4, which corresponds to ZT15. In contrast, in diurnal animals, assuming that GC peaks at ZT0 (*a* = π/2) and NE peaks at ZT18 (*b* = −π), both with an amplitude of 1, the resulting composite wave is as follows, making the peak *𝜙* =−π/4, which corresponds to ZT21.

To validate this model, we subsequently compared the timing of reported IOP rhythm peaks across different species. Unweighted meta-analysis revealed that diurnal animals, such as humans and monkeys, experienced peak IOP during the night(15–23), while nocturnal animals, including mice, rats, rabbits, and guinea pigs, also exhibited nocturnal elevation(12, 24–30). However, significant differences were observed between these two chronotypes, with peaks occurring at ZT15.43 in nocturnal and ZT20.75 in diurnal (*p* < 0.001) (Fig. 1E). These findings are similar to those presented in Fig. 1C,D, supporting the hypothesis that GC and NE independently and additively contribute to the circadian regulation of IOP.

### Combined framework can explain aging-modulated IOP rhythm changes in mice

To verify the validity of this model and its ability to explain variations in IOP rhythms, we next measured circadian IOP rhythms in young and extremely aged C57BL/6J mice under 12-h light/12-h dark cycles (Fig. 2A). The IOP was recorded every 4 h and fitted to a sine function to calculate its amplitude, acrophase, R^2^, and baseline (Fig. 2A). In aged mice, the IOP rhythm was altered (*p* < 0.0097 young vs. aged, two-way ANOVA) (Fig. 2B). When we analyzed the sine-curve-fitted individual data (Fig. 2C), IOP peak (acrophase) advanced by 1.77 h (*p* < 0.05, unpaired *t*-test), and the IOP level (baseline) also declined modestly (−0.64 mmHg; *p* < 0.01, unpaired *t*-test) (Fig. 2D). The amplitude tended to decrease to 80.7% of that in young controls (Fig. 2D). These changes indicate that aging reduces the strength of circadian IOP changes and shifts their timing earlier.

**Fig. 2.**
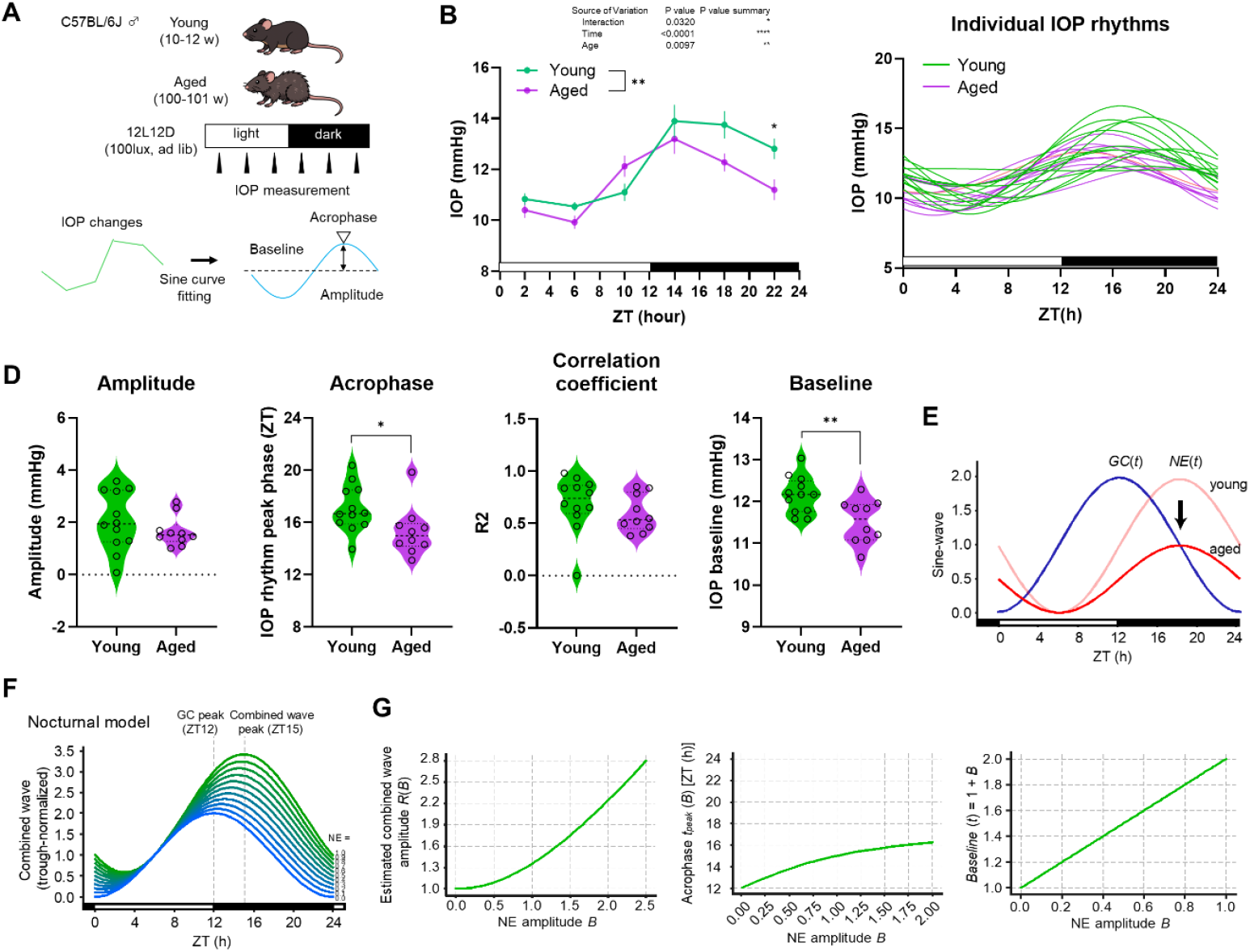
NE attenuation model explains aging-modulated IOP rhythm changes in mice. (A) Circadian IOP rhythms in young and extremely aged C57BL/6J mice. The animals were maintained under a 12-h light/12-h dark cycle, and IOP was measured every 4 h. Temporal profiles were fitted with a sine curve, from which the amplitude, acrophase, baseline, and goodness of fit (R^2^) were derived. (B) Circadian IOP rhythm in extremely aged mice. The phase advanced, and the amplitude decreased (^**^*p* < 0.01 young vs aged, two-way ANOVA with Šidák’s multiple comparison test). Data are shown as mean ± SEM (N = 10–12 animals). ZT0 denotes lights-on. (C) Sine curve fitting for individual mice. In aged mice, the amplitude tended to decrease, the phase significantly advanced by 1.77 h, and the baseline decreased by 0.64 mmHg (^*^p < 0.05, ^**^p < 0.01, unpaired t-test). (E) Phase and baseline changes in the additive waveform when the GC rhythm was fixed and only the amplitude of the NE rhythm was gradually reduced. (F) Relationship between NE amplitude and amplitude of the additive waveform. The phase advanced, and the overall amplitude decreased significantly. (G) Estimated amplitude and acrophase changes of the combined wave dependent on NE amplitude (*B*). Amplitude decreases and phase advances in the age model.

Age-related internal clock misalignments synchronize under LD conditions, constraining the cycle to 24 h and eliminating phase differences(32–35); therefore, they were excluded from this model. In humans, aging decreases most hormonal rhythm amplitudes, including melatonin, but minimally affects the GC rhythm(36). SCG-NE regulates the melatonin synthesis rhythm, and aging reduces the nocturnal melatonin peak but not the phase(37). By fixing the amplitude and phase of the GC and NE phases while varying only the NE amplitude, we examined how the combined IOP wave fluctuated (Fig. 2E). Assuming that in nocturnal animals, GC is fixed at the peak phase at ZT12 (*a* = −π /2) and NE at ZT18 (*b* = −π), and only amplitude B is reduced with amplitude A set to 1, the composite wave is given by this function (Fig. 2F):

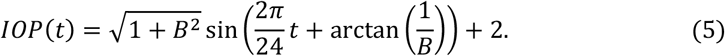

The amplitude function is as follows (Fig. 2G):

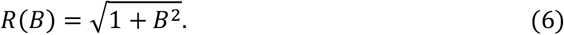

Therefore, the peak time *t*_*peak*_ (*B*) (ZT) of the composite wave is expressed as follows:

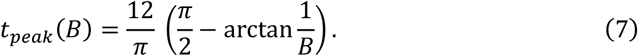

The *baseline (B*) function is as follows:

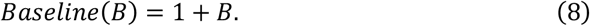

The reduction in NE amplitude advanced the phase and decreased the amplitude and baseline of the combined waveform (Fig. 3A–D). This composite wave model explains the variations in IOP rhythm with aging by changing the NE amplitude.

**Fig. 3.**
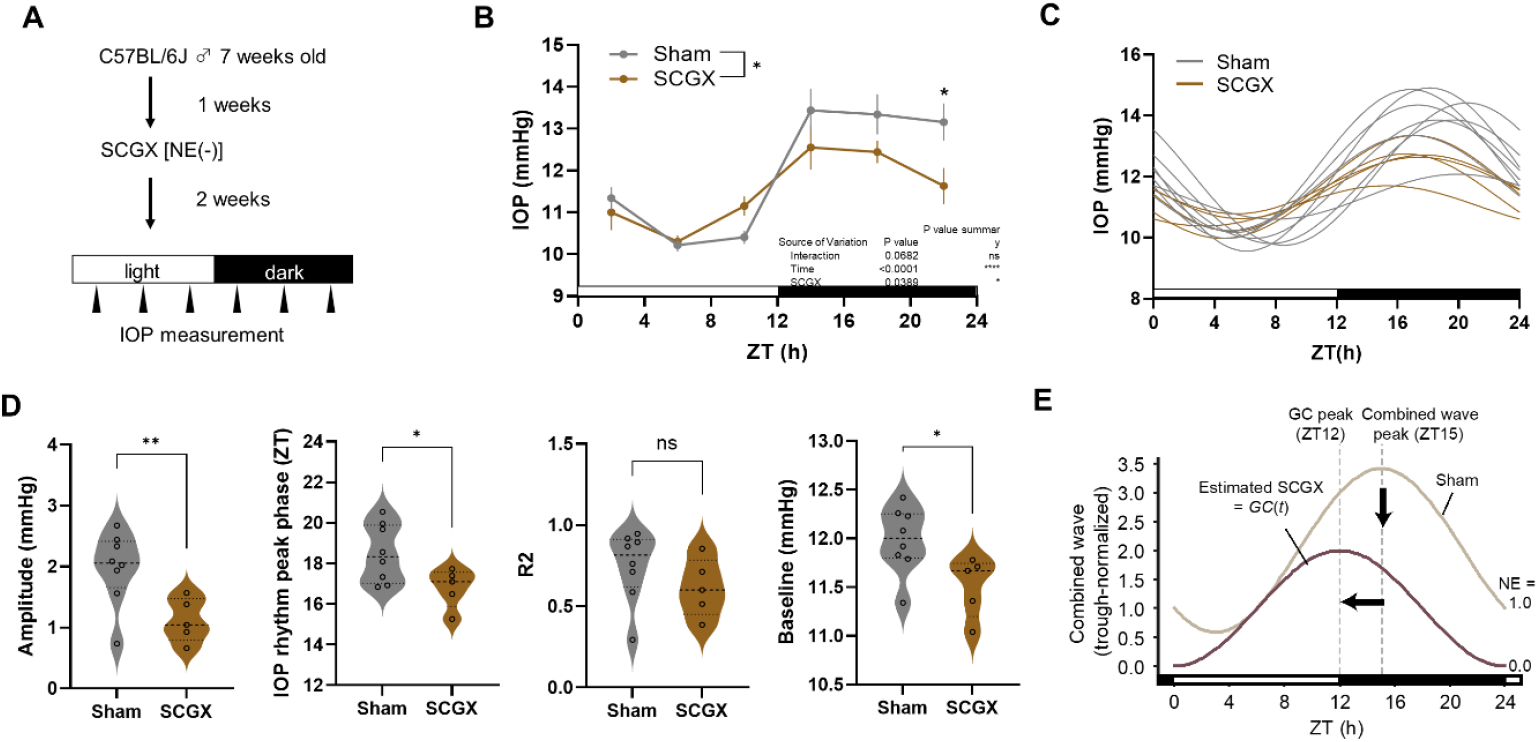
NE attenuation model interpretates effects of superior cervical ganglion (SCG) ablation on mouse IOP rhythms. (A) C57BL/6J mice were entrained under a 12-h light/12-h dark cycle for 1 week. Two weeks after SCG removal (SCGX) surgery, the circadian IOP was measured every 4 h. The sham- operated (N = 8) and SCGX (N = 5) groups were compared. (B) Circadian IOP rhythms in SCGX mice. SCGX attenuated the rhythm and induced phase advance (^*^*p* < 0.05 sham vs SCGX, two-way ANOVA with Šidák’s multiple comparison test). Data are shown as mean ± SEM (N = 5-8 animals). (C) Representative fitted sine wave. (D) Quantification of the amplitude, phase, correlation coefficient (R^2^), and baseline. Compared with the sham, SCGX reduced the amplitude by 50.1% and advanced the phase by 1.69 h, while left R^2^ was unchanged and the baseline was decreased (^*^*p* < 0.05, ^**^*p* < 0.01, unpaired *t*-test). These alterations closely resembled those observed in the aged mice. (E) Modeled combined waveform after SCGX, in which the NE component was eliminated, leaving only the GC rhythm.

### The NE attenuation model can show how SCG ablation is similar to aging effects

To confirm the contribution of sympathetic input, we performed bilateral SCGX in C57BL/6J mice. After two weeks, circadian IOP was measured every 4 h under light/dark cycles (Fig. 3A). SCGX altered IOP rhythm, similar to aging, compared to sham controls (*p* = 0.039, two-way ANOVA) (Fig. 3B). Analysis of the IOP waveforms (Fig. 3C) showed reduced amplitude by 50.1% (*p* < 0.01), advanced peak by 1.69 h (*p* < 0.05), decreased baseline pressure (*p* < 0.05), and unchanged curve fitting (R^2^) (Fig. 3D), paralleing the changes observed in aged mice (Fig. 2A–D). To simplify the model, eliminating NE input (amplitude → 0) advanced the phase by 3 h (*GC*(*t*)), qualitatively consistent with experimental data (Fig. 3E), thereby validating the effectiveness of this model. Although SGCX alters corticosterone rhythm in rats(38), its influence on GC rhythm cannot be ignored. Integrating these results showed that this combined wave captured IOP’s circadian characteristics (phase, amplitude, baseline) and helps understand IOP rhythm across species and aging.

### The NE attenuation model elucidates the effects of age on human IOP rhythm

Interestingly, in a human meta-analysis of age-related IOP rhythm changes (Fig. 4A)(5, 39–44), aging is not only associated with dumped IOP rhythm but also with delayed rather than advanced IOP rhythms (*p* < 0.05) (Fig. 4B-C). In contrast to the results in mice, IOP levels tended to increase with age (*p* = 0.05) (Fig. 4C). Assuming that in diurnal animals, the peak phase of GC is fixed at ZT0 (*a* = π/2) and NE at ZT18 (*b* = −π), and when amplitude A is set to 1 with only amplitude B being reduced as a variable (Fig. 4C), the resulting composite wave is given by the following function (Fig. 4E):

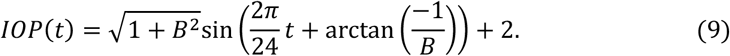

**Fig. 4.**
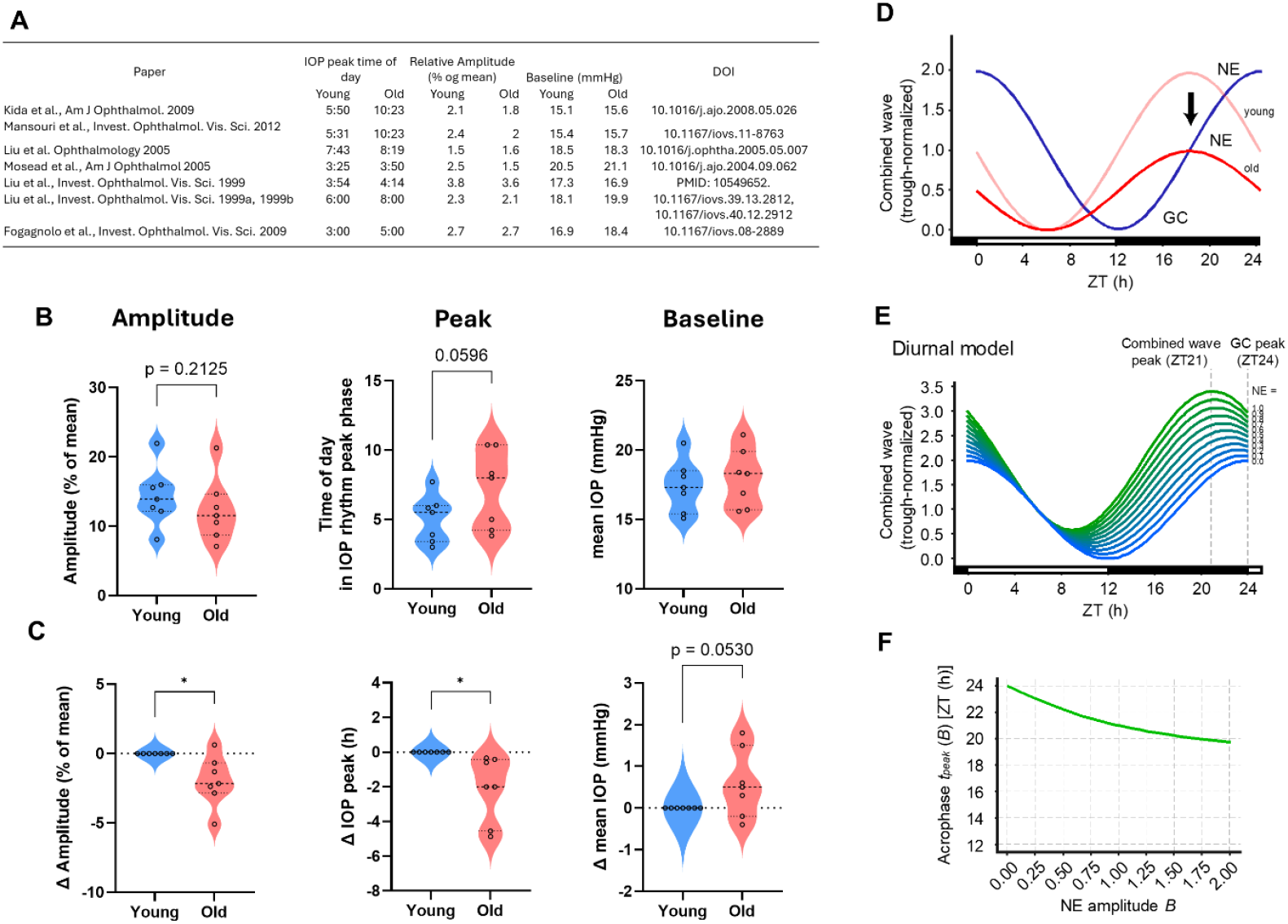
The NE attenuation model elucidates the effects of age on human IOP rhythm. (A) Table summarizing phase comparisons of circadian IOP rhythms between young and elderly humans. (B,C) Comparison of amplitude, peak, and mean of IOP rhythms between young and elderly humans and (C) normalized these differences. Aging was associated with a delayed IOP rhythm (*p* < 0.05, unpaired *t*-test). (D) Model of circadian IOP rhythm in diurnal humans. When the GC rhythm was fixed and the NE amplitude was reduced, the combined waveform showed a phase delay and decreased amplitude. (E) Estimated GC sine wave and halved NE sine wave in elderly participants. (F) Combined waveform in elderly humans showing a phase delay.

Although the amplitude and baseline (B) functions are the same as those of nocturnal animals, the peak time *t*_*peak*_ (*B*) is as follows (Fig. 4F):

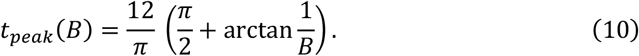

Our model predicted that reducing NE amplitude in the diurnal configuration (GC peak in the morning, NE at night) produces a phase delay rather than an advance (Fig. 3C– E), thereby accounting for the opposite direction of aging effects in nocturnal versus diurnal species by simply changing the amplitude of NE.

### Fluctuations in age-related GC rhythms do not affect the qualitative understanding of IOP rhythms in this model

To incorporate age-related GC changes into the model, we considered that while GC rhythm remains stable in middle-aged rats(45), aging in elderly humans and rodents causes rhythm attenuation through increased trough levels(46, 47). We constructed a model incorporating these age-related GC fluctuations:

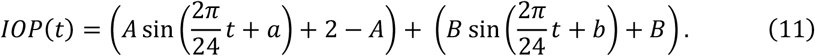

The phases were designated as *a* = –π/2 for nocturnal and +π/2 for diurnal, with b set to π (ZT18). Following previous studies (37, 46, 47), we assumed NE attenuation (50% amplitude, *B* = 0.5 [trough fixed]) and GC attenuation (70% amplitude, *A* = 0.7 [peak fixed]). The phases of the nocturnal and diurnal models shifted by 0.67 h (40 mins), with weaker phase fluctuations (SI Appendix, Fig. S1A,B). The baseline decreased from 2.0 to 1.8, reproducing the age-related IOP reduction. Since IOP levels were higher during the light phase, this parameter may account for age-related increases in human IOP only during this phase (Fig. 4). However, because this differs from the results in mice (Fig. 2), the NE attenuation model would likely be superior to the GC/NE attenuation model as a simple model.

## Discussion

Although IOP rhythms are established, their nocturnal rise in both diurnal and nocturnal species lacks a mechanistic explanation. We demonstrated that a model based on two sine waves representing adrenal GC and SCG-derived NE rhythms can account for species-, age-, and surgery-dependent IOP rhythm changes (Fig. 1). The direction of phase shifts and amplitude changes across diurnal and nocturnal animals and age groups could be explained by modifying the NE component’s amplitude while maintaining the GC component (Fig. 2, 4). Thus, the IOP rhythm represents the vector sum of two zeitgeber signals with distinct phases, rather than a single zeitgeber output. The model serves as a qualitative framework rather than a precise tool for predicting overall trends across various experimental conditions. While previous studies modeled circadian rhythms of peripheral tissues using single sinusoids, we propose that IOP rhythms emerge from multiple zeitgeber inputs with different phases.

Our meta-analysis findings from human studies qualitatively corroborate the GC-NE superposition framework predictions. Most studies have reported phase advancement and amplitude reduction with aging, aligning with model expectations. However, the results of baseline intraocular pressure (IOP) changes were less consistent. While age-related IOP increases are commonly reported in Western countries and are significant in the aging-glaucoma causal relationship(48, 49), East Asian countries like Japan and China show IOP decreasing with age(50, 51), as indicated by aged mice studies and mathematical models in the present study (Fig. 2). This suggests that regional differences are influenced by ethnicity, dietary habits, and climate. Large-scale global analyses are necessary to understand Western-type age-related IOP increases.

This study had several limitations. First, the model assumes a linear superposition of GC and NE, whereas actual interactions may include non-linear effects, receptor cross-talk, and saturation dynamics. Second, we treated GC and NE as fixed-period sinusoidal inputs, ignoring the variability introduced by light conditions, endogenous circadian rhythm patterns, feeding, and sleep–wake schedules. Third, the model does not incorporate aqueous humor production, outflow resistance, or ocular venous pressure, all of which contribute to the regulation of IOP. Fourth, differences in body posture, measurement devices, and cohort characteristics across studies limit the precision of the meta-analytic comparisons. Finally, the normalization procedure (setting trough = 0) simplifies the tonic components of the GC and NE signals, which may underestimate their baseline contributions.

This conceptual framework suggests promising clinical applications in the future. If IOP rhythms arise from GC and NE signal superposition, algorithms can determine component strength and phase using timed IOP readings with cortisol or autonomic markers. This could enable circadian IOP profiling to identify high-risk patients. The model suggests optimizing the therapeutic timing of β-blockers, prostaglandin analogs, or GC modulators based on the input phases. This framework offers a novel approach to time-based glaucoma management and chronotherapeutic scheduling.

Peripheral tissues receive multiple entrainment cues from feeding, temperature, melatonin, and hormones, suggesting broader implications of waveform superposition in circadian rhythms. Peripheral rhythms may reflect superimposed zeitgeber waveforms, providing insights into clock coordination and disease effects. Future studies should test this hypothesis by manipulating individual signals. Monitoring human cortisol and NE rhythms using IOP could reveal GC-NE contributions. Including feeding or melatonin rhythms may help establish a superposition model of peripheral regulation to inform therapeutic strategies.

## Methods

### Animals

Eight-week-old male C57BL/6JJmsSlc mice (Japan SLC Inc., Shizuoka, Japan) and 98-week-old male C57BL/6J-Aged mice (The Jackson Laboratory Japan, Inc.) were purchased and housed in plastic cages (215 × 316 × 150 mm) under a 12-h light (100 lx of fluorescent light)/dark cycle (12L12D, 0800 light ON, 2000 light OFF), and maintained at a constant temperature (24 ± 1 °C). Food (MF, Oriental Yeast, Tokyo, Japan) and water were provided *ad libitum*(31). All animal experiments were conducted in accordance with the Guidelines for the Faculty of Agriculture at Kyushu University. All experiments were approved by the Animal Care and Use Committee of Kyushu University (A23-223-2).

### IOP measurement

IOP measurements were performed using a tonometer (Icare TonoLab, TV02; Icare Finland Oy, Espmoo, Finland), as previously reported(7, 31). All mice were maintained under 12L:12D conditions for more than one week before IOP measurements. The unanesthetized mice were gently held using a sponge. IOPs were measured during the light phase under light (100 lx) conditions and during the dark phase under dim red light conditions. The IOP rhythm was obtained by measuring IOP at 4-h intervals under 12L12D conditions; at zeitgeber time (ZT) 2, 6, 10, 14, 18, and 22. ZT0 is the light onset time.

### SCGX

Superior cervical ganglionectomy (SCGX) was performed on 8-week-old male C57BL/6JJmsSlc mice under general anesthesia, as previously reported (52, 53). IOP measurements started 2 weeks after surgery for recovery.

### Analysis of rhythmicity

When statistically significant differences in IOP rhythm were observed by one-way analysis of variance (ANOVA), sine curve fitting was performed using GraphPad Prism 10 (GraphPad Software Inc., San Diego, CA, USA), as previously described (54).

### Mathematical modeling and curve fitting

Circadian intraocular pressure (IOP) rhythms were modeled as the superposition of two sinusoidal functions representing adrenal glucocorticoid (GC) and superior cervical ganglion–derived norepinephrine (NE) signals (1)-(4). Analytical derivations and illustrative simulations of the sine-wave model were performed in Python (v3.11) using SciPy (Virtanen et al., 2020), NumPy, and Matplotlib, with the code generated and debugged interactively in ChatGPT (OpenAI GPT-5). These simulations were used for conceptual validation, and experimental IOP data were independently fitted to sine curves using GraphPad Prism 10.

### Statistics and reproducibility

The results are shown as the mean ± standard error of the mean (SEM) of at least three independent experiments with five mice. Statistical comparisons were performed using GraphPad Prism 10 (GraphPad Software Inc., San Diego, CA, USA). Unpaired *t*-tests were used to compare two groups, and two-way ANOVA with Šidák’s or Dunnett’s multiple comparison test for more than three groups with two factors. Differences were considered statistically significant at *p* < 0.05.

## Data availability statement

All data are available from the corresponding author upon request.

## Acknowledgments

We are grateful to the Center for Advanced Technical and Educational Support, Faculty of Agriculture, Kyushu University. We would like to thank Editage (www.editage.com) for the English language editing. Additionally, we utilized the Paperpal tool to enhance the quality of the manuscript during its preparation. We express our gratitude to the Advanced Technology Support Center, Faculty of Agriculture, Kyushu University, for their support. This study was supported by The Hori Sciences and Arts Foundation, and the Mathematical and Data Science Education and Research Support Program of Kyushu University.

## Disclosure of AI usage

During the preparation of this work, the authors used ChatGPT (OpenAI GPT-5) and Paperpal to generate text, check grammar, develop a mathematical model, and assist with the literature search. After using this tool/service, the authors reviewed and edited the content as needed and took full responsibility for the content of the published work.

## Author contributions

K.I. was responsible for conceptualization, methodology, investigation, drafting of the original manuscript, and securing funding. R.F. and F.M. contributed to the development of the mathematical models. K.I., R.F., F.M., and S.Y. were involved in writing, reviewing, and editing the manuscript.

## Competing interests

The authors declare no conflicts of interest.

Correspondence and requests for materials should be addressed to Keisuke Ikegami.

## Supplemental Files

**SI Appendix, Fig. S1.**
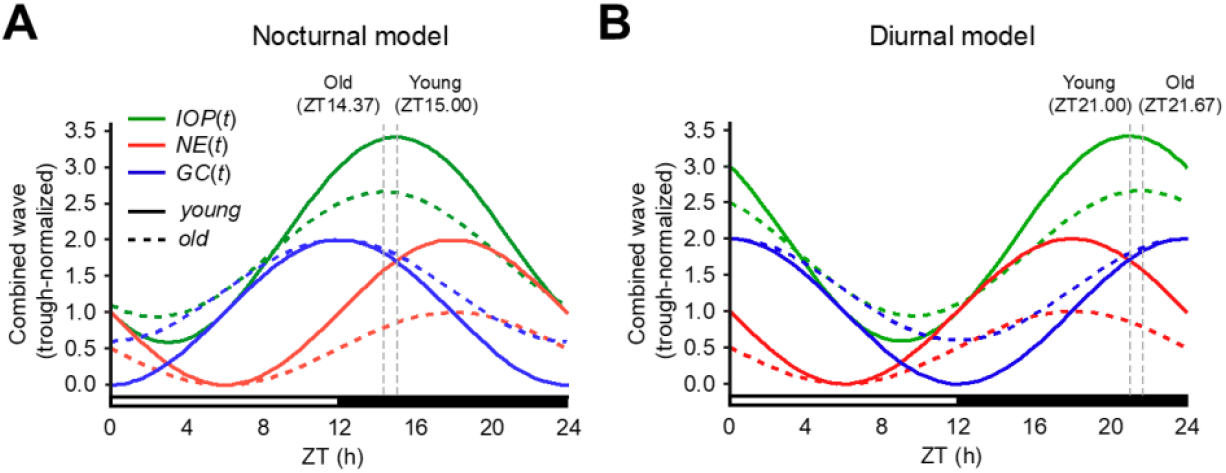
Alterations in age-related GC rhythms do not affect the qualitative understanding of IOP rhythms in this model. **(A)** IOP rhythm model in young and aged nocturnal animals. The IOP rhythm was modeled as the sum of two 24-h sinusoidal components representing GC (blue) and sympathetic NE (red): *a* = −π/2 (GC peak ZT12), *b* = π (NE peak ZT18), *A* = 0.7 (GC amplitude reduction with peak fixed), and *B* = 0.5 (NE amplitude reduction with trough fixed). The combined IOP waveform (green) peaked at ZT = 14.37, reproducing the slight phase advance observed in aged nocturnal species when the NE contribution declined. **(B)** IOP rhythm model of young and old diurnal animals. The IOP rhythm was modeled as the sum of two 24-h sinusoidal components representing GC (blue) and sympathetic NE (red): *a* =π/2 (GC peak ZT0), *b* = π (NE peak ZT18), *A* = 0.7 (GC amplitude reduction with peak fixed), and *B* = 0.5 (NE amplitude reduction with trough fixed). The combined IOP waveform (green) peaked at ZT = 21.67, reproducing the slight phase delay observed in aged nocturnal species when the NE contribution declined.

