## Supplemental Files for "Additive framework of hormonal waves explains species and age differences in circadian intraocular pressure rhythm"

**Classification line**

Biological Sciences | Physiology; Neuroscience; Computational Biology

**Keywords**

intraocular pressure rhythm; circadian biology; aging; glucocorticoid; norepinephrine; mathematical modeling; superior cervical ganglion; glaucoma

The authors declare that no conflicts of interest exist.

### Supplemental Files

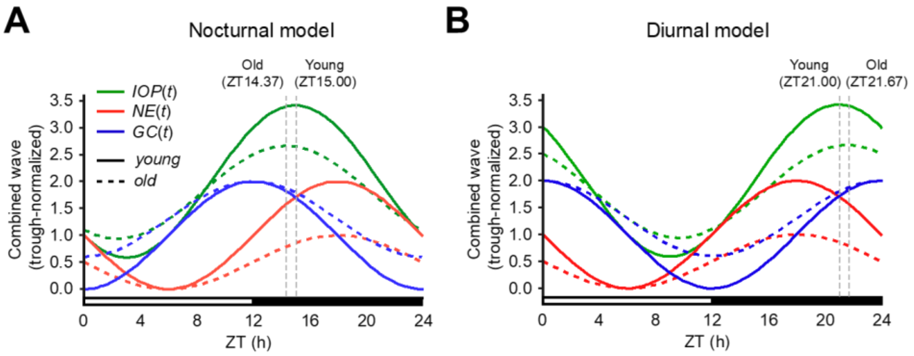


**SI Appendix, Fig. S1 Alterations in age-related GC rhythms do not affect the qualitative understanding of IOP rhythms in this model.**

**(A)** IOP rhythm model in young and aged nocturnal animals. The IOP rhythm was modeled as the sum of two 24-h sinusoidal components representing GC (blue) and sympathetic NE (red): $a = -\pi/2$ (GC peak ZT12), $b = \pi$ (NE peak ZT18), $A = 0.7$ (GC amplitude reduction with peak fixed), and $B = 0.5$ (NE amplitude reduction with trough fixed). The combined IOP waveform (green) peaked at ZT = 14.37, reproducing the slight phase advance observed in aged nocturnal species when the NE contribution declined. **(B)** IOP rhythm model of young and old diurnal animals. The IOP rhythm was modeled as the sum of two 24-h sinusoidal components representing GC (blue) and sympathetic NE (red): $a =\pi/2$ (GC peak ZT0), $b = \pi$ (NE peak ZT18), $A = 0.7$ (GC amplitude reduction with peak fixed), and $B = 0.5$ (NE amplitude reduction with trough fixed). The combined IOP waveform (green) peaked at ZT = 21.67, reproducing the slight phase delay observed in aged nocturnal species when the NE contribution declined.
